# Effects of acute intranasal allergen exposure on resident immune cells and sensory neurons in the mouse olfactory epithelium

**DOI:** 10.64898/2026.07.09.737488

**Authors:** Ryan E. Owens, Bailey E. Matthews, Michael A. Mastrangelo, Julian P. Meeks, Regina K. Rowe

## Abstract

The main olfactory epithelium (MOE) is the primary site of olfaction and consists of multiple cell types including olfactory sensory neurons (OSNs), sustentacular cells, and immune cells. Neuroimmune interactions in epithelial tissues are critical in maintaining tissue function, but how OSNs and immune cells interact in the MOE in healthy and diseased states is largely unknown. Cellular responses in the MOE determine how and whether OSNs maintain olfactory function and are repaired or replenished following inflammatory environmental exposures. We hypothesized that acute nasal aeroallergen exposure alters immune cell function in the MOE to elicit a neuroprotective response, thereby preserving OSN function. We developed an environmental aeroallergen exposure consisting of one week of daily intranasal house dust mite extract (HDM) instillations. Spectral flow cytometry indicated only subtle changes in resident immune cells proportions and phenotypes in the MOE. Immunohistochemical evaluation did not reveal extensive changes in immune cell distribution in the sensory epithelium or lamina propria, but instead we observed increases in axonal olfactory marker protein (OMP) expression in the lamina propria, where resident immune cells are most abundant. To evaluate the effects of HDM exposure on OSN function, we performed live *ex vivo* Ca^2+^ imaging of MOEs from HDM- and sham-exposed transgenic mice using objective-coupled planar illumination (OCPI) microscopy. OSN responses to multiple odorants revealed increased chemosensory sensitivity and decreased across-trial adaptation in HDM-treated epithelia. These results indicate that short-term nasal aeroallergen exposure minimally alters immune cell phenotypes, and instead induces functional changes in OSN physiology that preserve olfactory function.

## Introduction

The nasal epithelium is continuously exposed to foreign material, including microbes, aeroallergens, and industrial pollutants. The nasal epithelium houses the main olfactory epithelium (MOE), the principal site of olfaction (the sense of smell). Disruption of normal olfactory function can cause hyposmia (reduction in smell) or anosmia (complete loss of smell), which can significantly impact quality of life^1–4^. Inflammation is a common cause of olfactory dysfunction, and is induced by respiratory virus infections, bacterial sinusitis, allergic rhinitis, chronic rhinosinusitis, or direct tissue damage via exposure to environmental toxicants such as air pollution or smoking. Neuro-immune interactions are known to regulate olfactory function during significant inflammation^5, 6^, but the cellular responses to environmental inflammatory exposures are not well understood, creating a gap in our understanding of olfactory epithelial function.

The MOE consists of multiple cell types, including olfactory sensory neurons (OSNs), which detect odorous molecules, as well as supporting epithelial cells, secretory cells, and resident immune cells. Communication between neurons and immune cells has been investigated in the skin, gut, and the olfactory epithelium^5, 7–11^. Evidence suggests that bidirectional neuronal-immune communication is important in the MOE during states of severe inflammation and injury^12–15^, but few studies have investigated neuronal and immune responses to more common sources of nasal inflammation, for example repeated aeroallergen exposures. Given that many odorants enter the nasal passages alongside molecules with strong immunogenic properties (aeroallergens, for example pollen or dander), it is important to understand how neurons and immune cells respond in these mild, but extremely common sources of potential epithelial inflammation. Prior work revealed multiple changes in gene expression among neurons and immune cells in response to environmental exposure to odorants and aeroallergens^16^. In these studies, single-cell RNA seqencing identified changes in gene expression patterns in neuronal and resident immune cells in response to environmental exposure to a well-known aeroallergen, house dust mite (HDM) extract. Here, we report the development of an acute model of environmental aeroallergen exposure using daily intranasal HDM instillation, followed by functional measurements of resident immune cells and OSNs using high dimensional spectral flow cytometry, epifluorescence microscopy, and *ex vivo* volumetric live Ca^2+^ imaging (Figure 1). The results of these studies showed that acute HDM exposure induces subtle changes in immune cells, but causes significant changes to OSNs, including upregulation of axonal olfactory marker protein (OMP), increased odorant sensitivity, and decreased across-trial sensory adaptation. These results indicate acute environmental aeroallergen exposure modifies neuronal physiology without widespread reorganization of tissue immune cell populations, providing a foundation for future studies into MOE function during the many immune challenges that animals commonly encounter.

**Figure 1.**
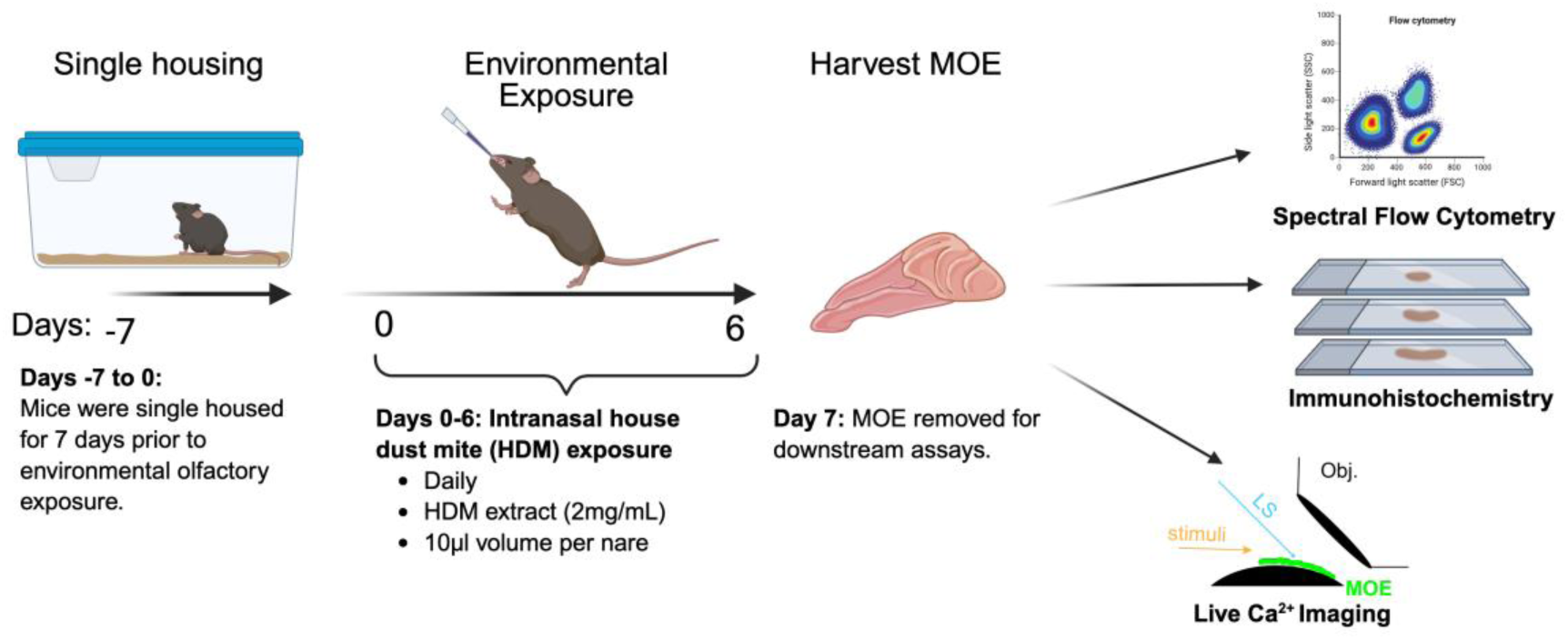
Experimental design for acute allergen exposure to HDM. Mice were single housed for 7 days prior to exposure. Mice received daily intranasal administrations of either house dust mite (HDM) extract (2mg/mL in PBS) or phosphate-buffered saline (PBS; sham control) for 7 consecutive days. Each mouse was administered 10 µL per nare (20 µL for 20µg total per animal, per day). One day following the final exposure, mice were perfused, and the main olfactory epithelium (MOE) was harvested for either immunohistochemistry, live *ex-vivo* Ca^2+^ imaging, or dissociated for spectral flow cytometry. MOE: main olfactory epithelium. LS: light sheet. Obj. imaging objective.

## Materials and methods

### Animals

All mice used in this study were originally purchased from Jackson Laboratories and used in accordance with the University of Rochester “University Committee on Animal Resources” (UCAR, the IACUC body for the university) under the guidelines of the National Institutes of Health. For spectral flow cytometry experiments, C57BL/6J mice (Strain #000664) were used. Transgenic mice were used for OSN physiology experiments. Specifically, mice expressing GCaMP6s in OSNs were generated by mating male Omp-cre^+/+^ mice, (B6;129P2- *Omp*^tm4(cre)Mom^/MomJ, JAX Stock #006668)^17^ to Ai96+/+ (Ai96(RCL-GCaMP6s, JAX Stock #024106)^18^ females, resulting in Omp^+/-^, Ai96^+/-^ offspring. All mice were used at 6-10 weeks of age. Figure legends indicate the number of male and female mice used in each experiment.

### Acute Allergen Intranasal Model

C57BL/6J mice (6-8 weeks, male and female) were single-housed for 7 days prior to environmental exposure (Figure 1). Mice then received daily intranasal administration of house dust mite (HDM) (Catalog #02.01.60, Citeq Biologics) extract daily for 7 days^19^. HDM was delivered to each nostril by pipetting 10μL of a 20 mg/mL solution (dissolved in Phosphate-Buffered Saline; PBS), under mild anesthesia. This volume preserved respiration without mechanical ventilation, and avoided deep HDM aspiration, which has the potential to induce lower airway distress. Sham control mice received the same volume of intranasal PBS (negative control). Following 7 days of repeated exposure, the MOE was dissected and either dissociated into a single-cell suspension as described below^16^ or processed for imaging. In a subset of experiments, one nasal hemisphere was processed by enzymatic digestion followed by antibody staining for spectral flow cytometry, and the other half was fixed in 4% paraformaldehyde (PFA) for imaging.

### Dissection and Dissociation of the MOE

Main olfactory epithelium (MOE) turbinates were dissected from the nasal cavity and transferred into freshly prepared cold oxygenated Ringer’s solution containing 115mM NaCl, 5mM KCl, 2mM MgCl_2_ hexahydrate, 2mM CaCl_2_ dihydrate, 25 mM NaHCO_3_, 10mM HEPES, and 10mM glucose with a pH of 7.4 at 4°C. Surrounding non-MOE tissue, including the septal organ and respiratory epithelium, was removed. Large bone and cartilage fragments were then removed without disrupting the epithelial layer under a dissection microscope (Leica Microsystems, Buffalo Grove, IL, USA). MOE tissue was subsequently transferred to 1.5 mL of digestion solution containing Dispase II (2 mg/mL, Sigma Aldrich Catalog #D4693), Collagenase Type 1 (2mg/mL, Sigma Aldrich Catalog #SCR103), Papain (0.25 mg/mL, Sigma Aldrich Catalog #10108014001), and DNase (2 mg/mL, Sigma Aldrich Catalog #DN25) in Ringer’s solution in a 6-well dish and minced under a dissection microscope. Tissue was incubated at 37 °C in a shaker incubator for 30–45 minutes, with intermittent mixing every 15 minutes. Following digestion, 5 mL of 37 °C Ringer’s-FBS (or Ringer’s-BSA) was added. Cells were gently triturated without introducing air bubbles and passed through a 40-μm filter (gravity facilitated) into a 50 mL conical tube. The culture dish and filter were rinsed with an additional 10 mL of Ringer’s-FBS to collect remaining cells. Cells were kept on ice and centrifuged in a swinging-bucket rotor at 300 × g for 7 minutes at 8 °C. The supernatant was removed, and the pellet was resuspended in 10 mL of Ringer’s solution. The suspension was triturated using a 200-μL pipette tip, passed through a 40-μm filter to remove remaining debris, and centrifuged at 300 × g for 5 minutes at 4 °C. The final pellet was resuspended in 500 μL of PBS. Cell viability was subsequently assessed using trypan blue, with a target viability of >80%. Murine spleens were processed in parallel as a positive control for each antibody target used for flow cytometry.

### Spectral flow cytometry

Following dissociation, cells were first stained for viability with Live-Dead Blue per manufacturer’s instructions. Cells were then incubated in conjugated monoclonal antibodies (Supp. Table 1) for 45 min at 4°C and washed with PBS before being fixed in 2% formalin. Following formalin fixation, cells were re-suspended in 150 μL of PBS before cell events were acquired on a Cytek Aurora spectral flow cytometer (Cytek Biosciences, Fremont, CA). Live spectral unmixing was performed during acquisition by the Cytek SpectroFlo software using single fluorochrome stained beads for each antibody (UltraComp Compensation Beads, ThermoFisher Scientific), a live/dead single stain cell control, and an autofluorescence extraction using unstained MOE cells. Unmixed data was analyzed using FlowJo^TM^ version 10.10 (BD Life Biosciences) followed by statistical analysis in Prism GraphPad Version 11.

**Table 1:** Olfactory stimulus panel. MOE tissue was exposed to each stimulus 4x, with each stimulus (odorant, concentration) lasting 15 seconds. Female mouse urine (FMU) was used at 1:100 as a positive control. Odors from four different functional groups were used: amyl acetate (AA, ester), octanal (Oct, aldehyde); acetophenone (Acet, aromatic ketone), and 1,8 cineole (a.k.a. eucalyptol, Euc, alkane). Ringer’s solution was delivered for 30s between each stimulus trial to flush away prior stimuli, and 0.1% DMO in Ringer’s saline used as a negative control stimulus.

| Olfactory stimuli | Concentrations |
| --- | --- |
| Female mouse urine (FMU) | 1:100 |
| Amyl Acetate (AA) | 100μM, 30μM, 3μM |
| Acetophenone (Acet) | 100μM, 30μM, 3μM |
| Octanal (Oct) | 100μM, 30μM, 3μM |
| 1,8 Cineole (Euc) | 100μM, 30μM, 3μM |
| DMSO (Vehicle) | 0.1% |

### Spectral flow cytometry analysis

We used a multi-step gating strategy to identify major immune cell populations and subsets using a panel of well characterized antibodies (Supp. Fig. 1). Immune cell subsets were determined from events that were further gated based on specific combinations of surface protein expression. Major immune cell populations were identified from live, single, CD45^+^ cells. Gating of immune cell populations were as following: DCs (CD11c^+^, MHC II^+^), Neutrophils (CD11b^+^, Ly6G^+^), Eosinophils (CD11b^+^, Ly6G^HI^, Siglec-F^HI^); CD8^+^ T cell (CD3^+^, CD8a^+^); CD4^+^ T cells (CD3^+^, CD4^+^), B cells (CD19^+^, CD3-) and Macrophages (CD11b^+^, F4/80^+^). Monocytes were CD11b^+^F4/80^low^ and then separated from Ly6C high and low populations. Macrophage subsets were identified as CX3CR1^+^ macrophages (MHCII^hi^, CX3CR1^+^ or CX3CR1^+^ MHC II^low^); and NK cells (NK1.1). FlowJo^TM^ Software v10.10 (BD Life Sciences) was used for analysis. Single-color controls as well as stained control spleen tissue were used to confirm gating of immune cell populations.

### Immunohistochemistry

C57BL/6J mouse MOEs were sectioned and stained to evaluate the spatial localization of CD45^+^ immune cells and OMP^+^ neurons following HDM exposure or sham controls. Adult mice under deep ketamine/xylazine anesthesia were exsanguinated using ice-cold phosphate buffer (PB), followed by transcardial perfusion 4% paraformaldehyde in PB. The snout and olfactory bulb were dissected and post-fixed for 24-48 hours in 4% paraformaldehyde-PB. In a subset of experiments, a single hemisphere of mouse snouts used for acute MOE dissociation (described above) were post-fixed in 4% paraformaldehyde-PB for 1 week. Following fixation, samples were decalcified in 10% ethylenediaminetetraacetic acid (EDTA; pH 7.4) in water for 1 week. Finally, the tissues were cryoprotected in high-sucrose PB (20%, 30%) before embedding in Optimal Cutting Temperature compound (OCT). Sections were cut at a thickness of 25-*μ*m using a Leica CM1850 cryostat (Leica Biosystem) and mounted onto poly-L-Lysine coated adhesive slides and stored at -20°C. Slides were baked at 65°C for 2 hours, then lowered to room temperature (RT) for 15 min. Slides were then washed with 1X PBS (3 times for 5 mins each), then washed with PBS containing 0.1% Triton-x (for 5 mins) followed by blocking with a 2-3 h incubation in PBS with 0.1% Triton X and 10% normal goat serum (NGS). Proteins were stained with primary antibodies for OMP (1:1000) and CD45 (1:250) at 4°C in PBS containing 0.1% Triton X and 1% NGS. Primary antibodies were washed (3 times for 5 mins each) in PBS containing 1% Triton X and 1% NGS at RT. Secondary antibodies were diluted in PBS containing 0.1% Triton X and 1% NGS (Goat anti-rabbit; 1:1000), (Goat anti-rat 488; 1:500). Nuclear DAPI staining was used to identify cell nuclei (1:1000).

### Live ex-vivo imaging

#### Olfactory stimulus presentation

All olfactory stimuli were purchased from Sigma-Aldrich (Table 1). The panel of stimuli was crafted to represent compounds from diverse functional groups with well-characterized olfactory properties. Individual stimuli, including amyl acetate (ester), eucalyptol (1,8-cineole), octanal (aldehyde), and acetophenone (aromatic ketone), were initially prepared as 100 mM stock solutions and dissolved in 0.1% dimethyl sulfoxide (DMSO) (Table 1). These stock solutions were serially diluted in Ringer’s solution to final concentrations of 100 µM, 30 µM, and 3 µM. Female mouse urine (FMU) was collected from C57BL/6J female mice aged 6 to 8 weeks. Briefly, urine was collected freshly by flash-freezing in liquid nitrogen, stored at −80°C, thawed, pooled across individual mice, and sterile filtered. FMU was diluted at 1:100 in control Ringer’s solution. All Ringer’s solutions and working dilutions were prepared fresh on the day of imaging.

#### Intracellular Ca^2+^ Imaging

Imaging of MOE tissue was performed on a custom Objective Coupled Planar Illumination (OCPI) microscope adapted from Holekamp et al., 2008^20^. Illumination was at 488 nm, the light sheet was ∼5 μm thick, and the objective was a liquid-immersion 20x, 0.5 NA (Olympus). The camera was a pco.edge 4.2 sCMOS (PCO-Tech/Excelitas). Tissue was placed on nitrocellulose paper, and a custom 3D printed chamber clip was used to hold the tissue in place and facilitate direct application of stimuli. Volumetric imaging produced datasets encompassing 615 x 615 x 200-400 μm, with each volume consisting of 51 frames at 1024 x 1024 original resolution. The exposure time for each stimulus frame was 50 ms (∼2.5 s per volume), and the inter-stack interval was set at 3 s (0.33 Hz imaging). Each stimulus was presented for 5 stacks (∼15s), with inter-stimulus intervals set at 10 stacks (∼30s). Postprocessing reduced each volume to 513 x 513 x 51 voxels prior to image registration. A minimum of 4 repeats of each stimulus (odorant, concentration) was delivered to each tissue.

#### Data Analysis of volumetric MOE Ca^2+^ imaging

Data analysis was performed using MATLAB software. Neuronal responses were recorded and underwent rigid registration before the 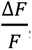, the relative change in GCaMP6s/f intensity, was calculated by subtracting the mean pre-stimulus intensity from the mean of the first 3 stacks (9 s) during test stimulus delivery, then dividing by the value of the mean pre-stimulus delivery. Volumetric regions of interest (ROIs) were manually drawn around the cell bodies of well-registered OSNs that reliably responded to stimulation in two or more repeats to exclude spontaneously active neurons, and to ensure inclusion of OSNs that experienced stimulus-dependent sensory adaptation. Following ROI selection, the mean intensity for each ROI was calculated for all image stacks, including each stimulus, concentration, and trial, to generate a matrix of fluorescence intensities. Stimulus responsiveness was determined by comparing the across-trial stimulus-evoked 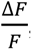, of each ROI to the across-trial negative control stimuli (0.1% DMSO in Ringer’s saline). ROIs demonstrating an across-trial p-value below 0.2 and mean initial 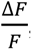 response to one or more test stimuli exceeding 10% were further analyzed. An unpaired, two-tailed Student’s *t*-test was used to assign p-values to screen initial responsiveness. Subsequently, nonparametric tests (Wilcoxon-Mann-Whitney) were used to assign p-values, and nonparametric estimates of effect sizes (Hodges-Lehmann estimator) and bootstrap 95% confidence intervals (1,000 bootstraps) were calculated using custom MATLAB software.

## Results

### Assessing MOE immune cell abundance following HDM treatment

In previous studies, we observed gene expression changes in OSNs and MOE resident immune cells following one week of environmental HDM exposure, suggesting that HDM treatment alters neuro-immune interactions in the MOE^16^. We adapted this acute HDM paradigm to directly introduce HDM into the nasal cavity (Fig. 1, see Materials and Methods). Wild type male and female C57BL/6J mice were intranasally exposed to HDM daily for 7 days, with sham animals receiving intranasal PBS instillation. One day after the final exposure, MOEs were dissociated into single cell suspension, and immune cell populations were identified by spectral flow cytometry (Fig. 2, Supp. Table 1). This flow cytometry panel successfully identified the major immune cell populations and subsets within the MOE (Fig. 2, Supp. Fig. 1). Comparisons between different treatments (HDM vs Sham) were performed using unpaired, two-tailed Student’s *t*-test assuming a Gaussian distribution and equal standard deviations. The total proportion of CD45^+^ cells was not significantly increased in HDM-treated mice compared to sham-exposed mice (Fig. 2A). The proportion of CD45+ cells that were eosinophils, neutrophils, macrophages, and monocytes were also unchanged in the HDM-treated group compared to sham (Fig. 2D-E). We observed an increase in dendritic cells (DCs, Fig. 2F, p = 0.04) and natural killer (NK) cells (Fig. 2G, p = 0.01) in the MOE of HDM-exposed mice compared to sham. Among the T cell (CD3^+^) populations, we identified no significant difference in the percentage of either CD4^+^ or CD8^+^ T cells (Figure 2I-J). In other epithelial tissues, CX3CR1^+^ macrophages have been associated with surveillance in gut and skin tissues^21, 22^. We evaluated CX3CR1^+^ cells in the MOE response to HDM exposure and observed no change in this population following HDM treatment (Supplementary Fig. 2). Collectively, flow cytometry analysis indicates a minimal/low-grade inflammatory response in the MOE upon HDM exposure, indicating this treatment does not cause widespread inflammation, infiltration, or reorganization of immune cell populations.

**Figure 2:**
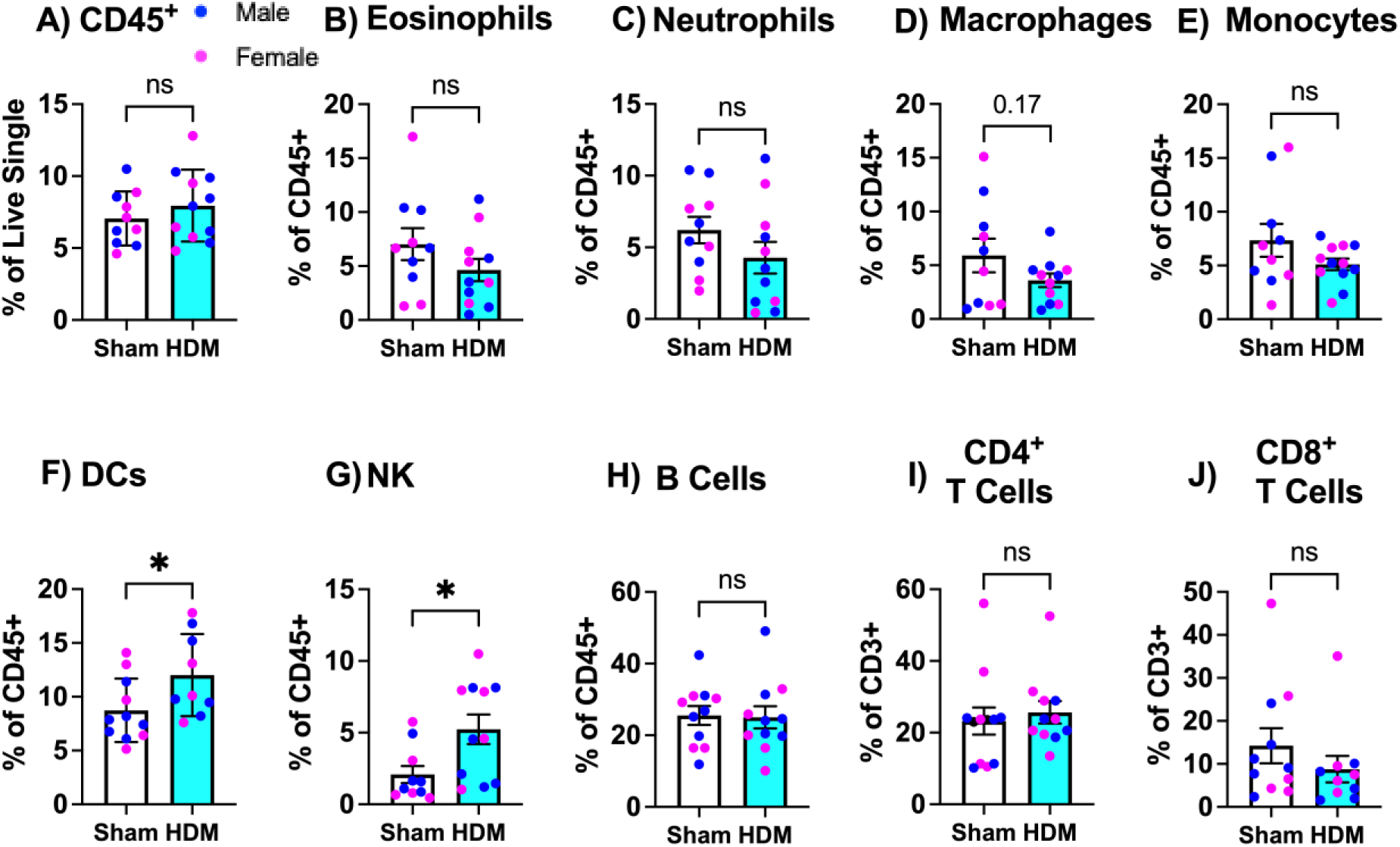
Evaluation of major immune cell subsets following acute HDM exposure. Wild-type C5BL/6J mice (n=11) were exposed to either Sham (white) or HDM (teal) intranasally for 7 days. MOEs were dissociated and flow cytometry performed for (**A**) total CD45^+^ immune cells as a percent of live singles, and major immune cell subsets (**B-H**) as a percent of CD45^+^ population, or (**I-J**) T cell subsets as a % of total CD3^+^ T cells. In each graph, magenta symbols indicate results for females, and dark blue symbols indicate results for males. * p < 0.05 (unpaired, two-tailed Students *t*-test); ns: non-significant.

### Regional variability in MOE immune cell abundance

The CD45+ immune cell population in the MOE includes diverse cell types (Fig. 2)^23–25^. These cells are not uniformly distributed within the MOE, but instead primarily reside in the lamina propria, with rarer cells resembling microglia localized in the apical epithelium, intermingled with OSNs (Supp. Fig. 3). The distribution of CD45+ cells also varies between anatomical subdivisions of the MOE (i.e., across the anterior-dorsal to posterior-lateral axis)^26–29^. We therefore investigated potential differences in the abundance or identity of immune cell populations across MOE zones (Fig. 3). In these experiments anterior/dorsal turbinates were dissected and isolated separately from the posterior-lateral turbinates. Using the same flow cytometry panel (Fig. 2, Supp. Fig. 1, Supp. Table 1), we investigated potential differences in immune cell populations in different MOE regions. Comparisons across two different tissue regions were made using an unpaired, non-parametric Mann-Whitney U test. The overall abundance of CD45^+^ immune cells was increased in anterior MOE samples (Fig. 3A, “Turb 1”) compared to more posterior-lateral turbinates (Fig. 3A, “Turb 2-6”, p= 0.004). In addition to a difference in the total number of CD45+ cells in the anterior-dorsal MOE, this comparison also identified differences in the proportion of specific immune cell subsets (Fig. 3B-J). We identified a relative increase in the abundance of eosinophils in the posterior (Turb 2-6) region compared to anterior/dorsal region (Turb 1, p = 0.004, Fig. 4B). This suggests that the increase in total CD45^+^ populations in the anterior/dorsal region is not due to an increase in eosinophils in this region. We also observed a relative decrease in the abundance of B cells in the posterior region (Turb 2-6) compared to the anterior region (Turb 1, p =0.001, Fig. 3H). The increased abundance of B cells could be result of an association between B cells and Bowman’s glands, which are more plentiful in the anterior/dorsal region^30–35^. Finally, we observed a small increase in the proportion of CD4^+^ T cells in the posterior/lateral region (Turb 2-6) compared to the anterior region (Turb 1, p = 0.03, Fig. 3J). Additional tests of regional differences in macrophage and monocyte populations revealed that MHC II-high macrophages were enriched in the posterior turbinates compared to anterior turbinates (Supp. Fig. 4). Similarly, Ly6c-low monocytes were enriched in the posterior turbinates (Supp. Fig. 4). Overall, these results show that the MOE has regional differences in immune cell distributions that may affect the response to environmental exposure to immunostimulatory molecules, including HDM.

**Figure 3:**
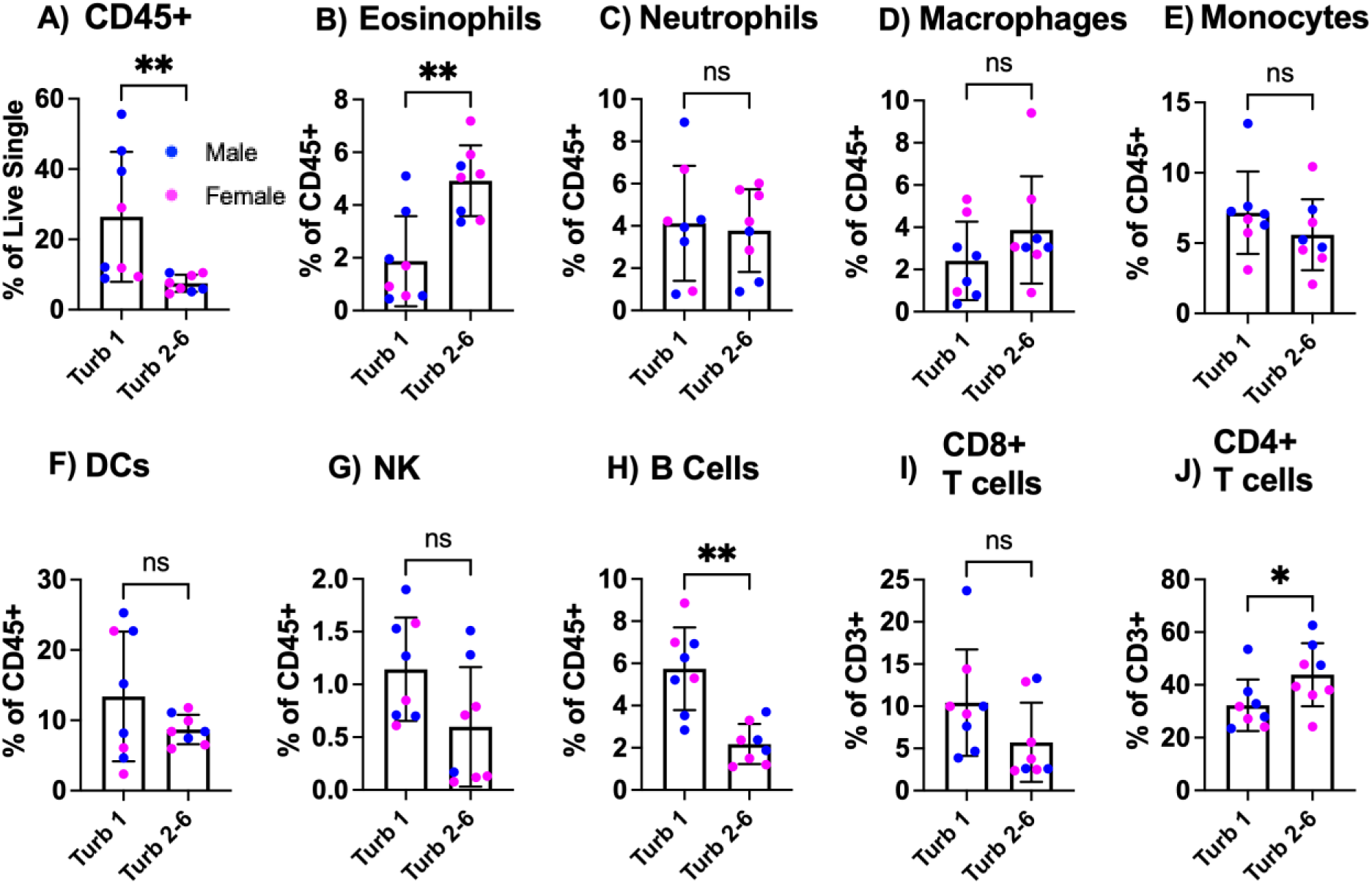
Immune cell distribution across MOE regions. MOEs from in C57/BL6J mice (n=8) were isolated and separated into anterior/turbinate 1 (Turb 1) and the more posterior-lateral turbinates (Turb 2-6). Tissue was dissociated and flow cytometry performed as in Fig. 2. Turb 1 vs. Turb 2-6 were compared for **(A)** CD45^+^ cells as a percent of live single events, **(B-H)** main immune cell subsets as a percent of total CD45^+^ cells, and **(I-J)** CD8^+^ and CD4^+^ T cell subsets as a percent of total CD3^+^ T cells. In each graph, magenta symbols indicate results for females, and dark blue symbols indicate results for males. * p < 0.05 (Mann-Whitney test); ** p < 0.01; ns: non-significant.

**Figure 4.**
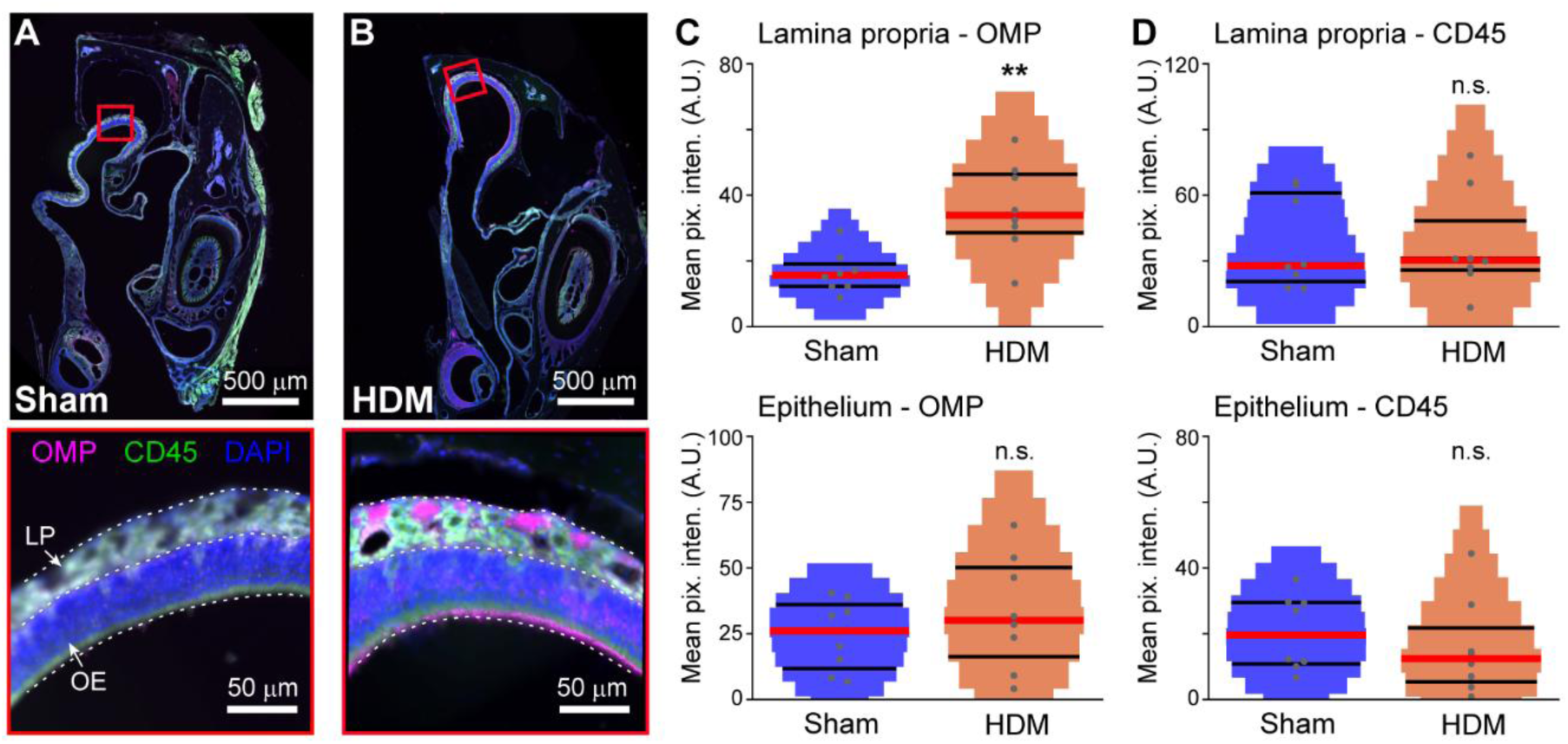
HDM exposure alters axonal olfactory marker protein expression. (**A-B**) Micrograph of anterior/dorsal portions of the olfactory epithelium of sham (**A**) and HDM-treated (**B**) mice. Insets at bottom highlight olfactory epithelial (OE) layer and lamina propria (LP), including bundles of OSN axons. OMP: olfactory marker protein (magenta). CD45 (green). DAPI (blue). (**C**) Mean OMP-labeled pixel intensity in the lamina propria (top) and epithelial layer (bottom). ** p < 0.01, Kruskal-Wallis test. Hodges-Lehmann estimate of effect size: 18.3 units, bootstrap confidence interval (1,000 bootstraps): [9.54 – 31.26]. n.s. not significant (p > 0.05). (**D**) Mean CD45-labeled pixel intensity in the lamina propria (top) and epithelial layer (bottom).

### OMP increases in OSNs following HDM exposure

Even though the overall abundance and relative distributions of immune cells were minimally changed by HDM treatment (Fig. 2), flow cytometry cannot assess potential changes in their spatial localization within the epithelium. Moreover, these experiments did not evaluate OSNs themselves, which also have altered patterns of gene expression following HDM exposure^16^. We therefore evaluated the spatial distribution of immune cells in the MOE relative to OSNs (Fig. 4). Because of the observed regional differences in immune cell distributions, we focused our evaluation on the anterior/dorsal region, which had a higher overall abundance of CD45+ cells, and resides nearest the nostrils, where the HDM treatment is introduced. We performed immunohistochemistry staining for OMP protein, a traditional marker of mature OSNs and cAMP buffer^36^, and for CD45, a broad marker of immune cells, in the anterior/dorsal MOE. We observed robust CD45 staining in the epithelial layer and lamina propria (Fig. 4A-B). We quantified OMP and CD45 pixel intensity in both the epithelial layer and the lamina propria (Fig. 4C-D). Consistent with our flow cytometry, we saw no indication of a change in overall CD45 intensity in either the olfactory epithelium or lamina propria (Fig. 4D). In contrast, we did observe an increase in OMP staining intensity, specifically in axon bundles in the lamina propria of HDM-treated animals (Fig. 4C). Alterations of OMP expression in axons (and olfactory bulb glomeruli) have been associated with olfactory tissue expansion and remodeling in models of olfactory dysfunction^12, 37–42^, suggesting that HDM treatment had a similar effect.

### HDM alters OSN odorant sensitivity

The changes in OMP expression in the anterior/dorsal MOE, along with prior evidence that HDM treatment alters OSN gene expression patterns^16^, suggested potential effects on OSN physiology. OMP disruption does not cause anosmia but does cause changes in olfactory function^36, 42–45^. We therefore tested whether acute HDM exposure results in changes to OSN physiology. We performed acute intranasal HDM exposure in Omp-cre^46^ x Ai96^17^ mice, which express the genetically encoded Ca^2+^ indicator GCaMP6s in the cytoplasm of mature OSNs. Following the 7-day HDM exposure paradigm, the anterior/dorsal MOE was harvested and subjected to live *ex vivo* imaging with OCPI microscopy, a light sheet-based volumetric imaging method. This approach allowed us to measure the Ca^2+^ responses of hundreds to thousands of OSN per tissue to multiple stimulus trials of a panel of well-characterized odorants at multiple concentrations (Table 1, Fig. 5A-C). Individual OSNs demonstrated combinatorial, concentration-dependent responses to stimulation (Fig. 5D), consistent with previous studies in the MOE^47^ and vomeronasal organ^48^. Across 21 MOE tissues from 16 mice in both HDM-treated and sham controls, and across both sexes, we observed OSNs with functionally similar neural response profiles, which clustered into 7 groups (Fig. 5E-F). Clusters included prominent, broadly tuned OSNs (Fig. 5E, Clusters C2 and narrowly tuned OSNs (Fig. 5E-F, Clusters 1, 3, 5-7). The presence of functionally distinguishable groups is consistent with the presence of OSN populations that express either broadly or narrowly tuned odorant receptors in the anterior/dorsal MOE^47^.

**Figure 5.**
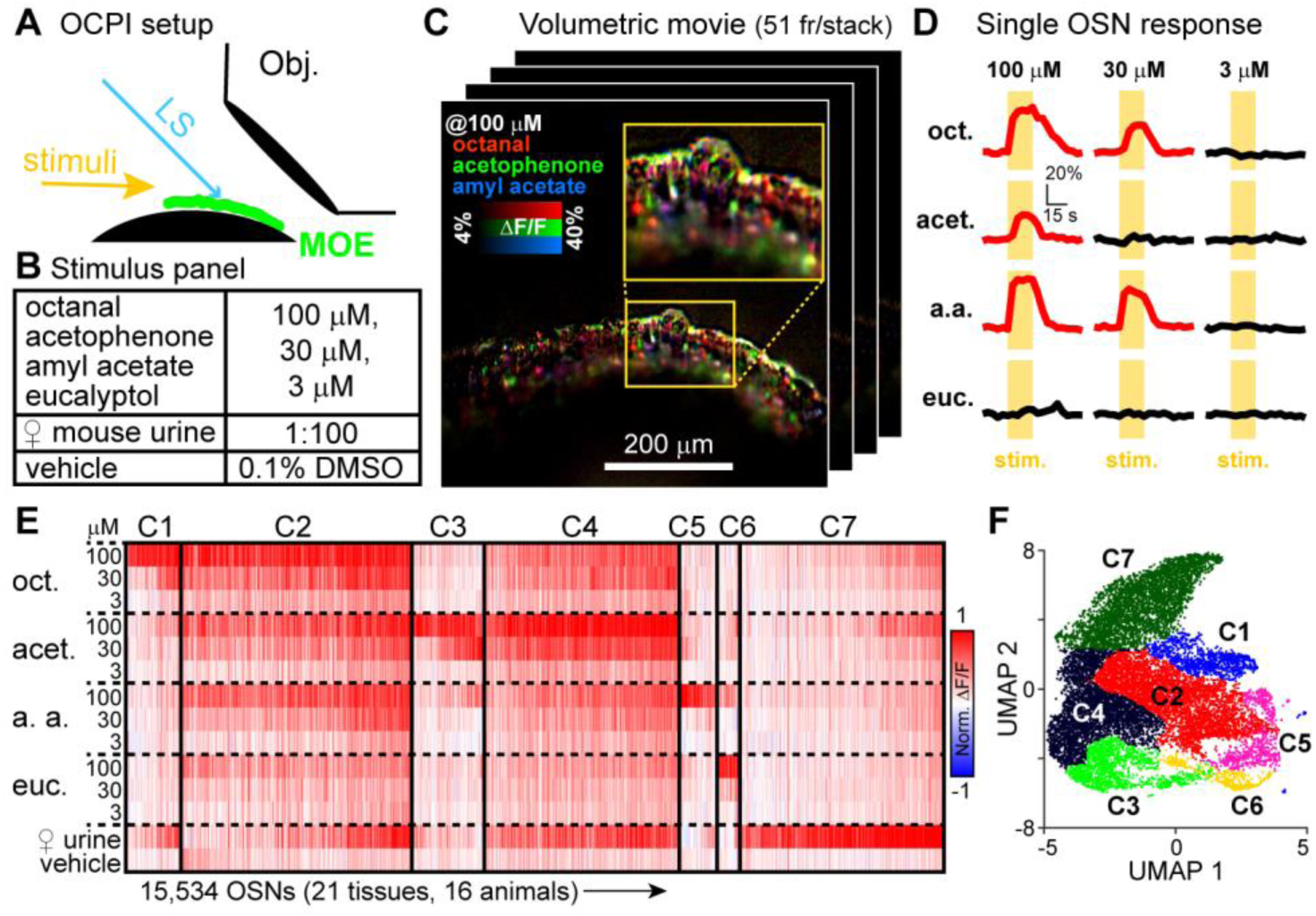
Population OSN GCaMP6s Ca^2+^ imaging via objective-coupled planar illumination (OCPI) microscopy. (**A**) OCPI setup. LS: light sheet. Obj.: imaging objective. (**B**) Stimulus panel used in study. Each stimulus was applied to the tissue 4 times in a randomized, interleaved block design. (**C**) Colorized image of average ΔF/F responses in an example tissue. Each pixel in the image was colorized according to its responsiveness to independent stimulation with 100 μM octanol (red), acetophenone (green), and amyl acetate (blue). Color mixtures (yellow, white, etc.) indicate responses to multiple ligands. Responsive OSN cell bodies are clearly visible and were selected as regions of interest (ROIs) for subsequent analysis. (**D**) Example OSN across-trial average ΔF/F responses to each monomolecular odorant at each concentration. Yellow rectangles indicate the stimulus application timing. (**E**) Clustered heatmap of normalized ΔF/F responses for 15,534 OSNs obtained from 21 tissues (16 animals). Cluster boundaries are indicated by solid vertical dividers. C1-C7 refer to clusters of functionally similar of OSNs, further illustrated in Panel F. (**F**) Uniform manifold approximation and projection (UMAP) plot of OSN functional clusters. C1-C7 refer to the clusters identified in Panel E.

We first used OCPI imaging data to evaluate the sensory tuning properties of individual OSNs and populations of OSNs (Fig. 6). Each of the 4 odorants in the stimulus panel was delivered to the tissue at 3 pre-determined concentrations, 3, 30, and 100 μM in saline solution, and at least 4 independent trials of each odorant were delivered to each tissue (Fig. 6A). Each OSN’s across-trial mean responses to each odorant were fit with the Hill Equation to identify each cell’s EC_50_ value (concentration eliciting normalized ΔF/F values representing 50% the maximum response, Fig. 6B). This allowed us to directly test whether HDM-exposed tissues experienced a shift in the sensitivity to odorants (Fig. 6C). The results of this analysis indicated a leftward-shift in the EC_50_ value of OSNs in HDM-exposed tissues compared to sham controls (Fig. 6B-F). This shift was observed for multiple odorants, suggesting that the effects were not restricted to a single odor-receptor pair (Fig. 6D-F). These results indicate that HDM-exposed OSNs demonstrate higher odorant sensitivity, and imply that HDM treatment increases, rather than decreases, OSN responses to odorants.

**Figure 6.**
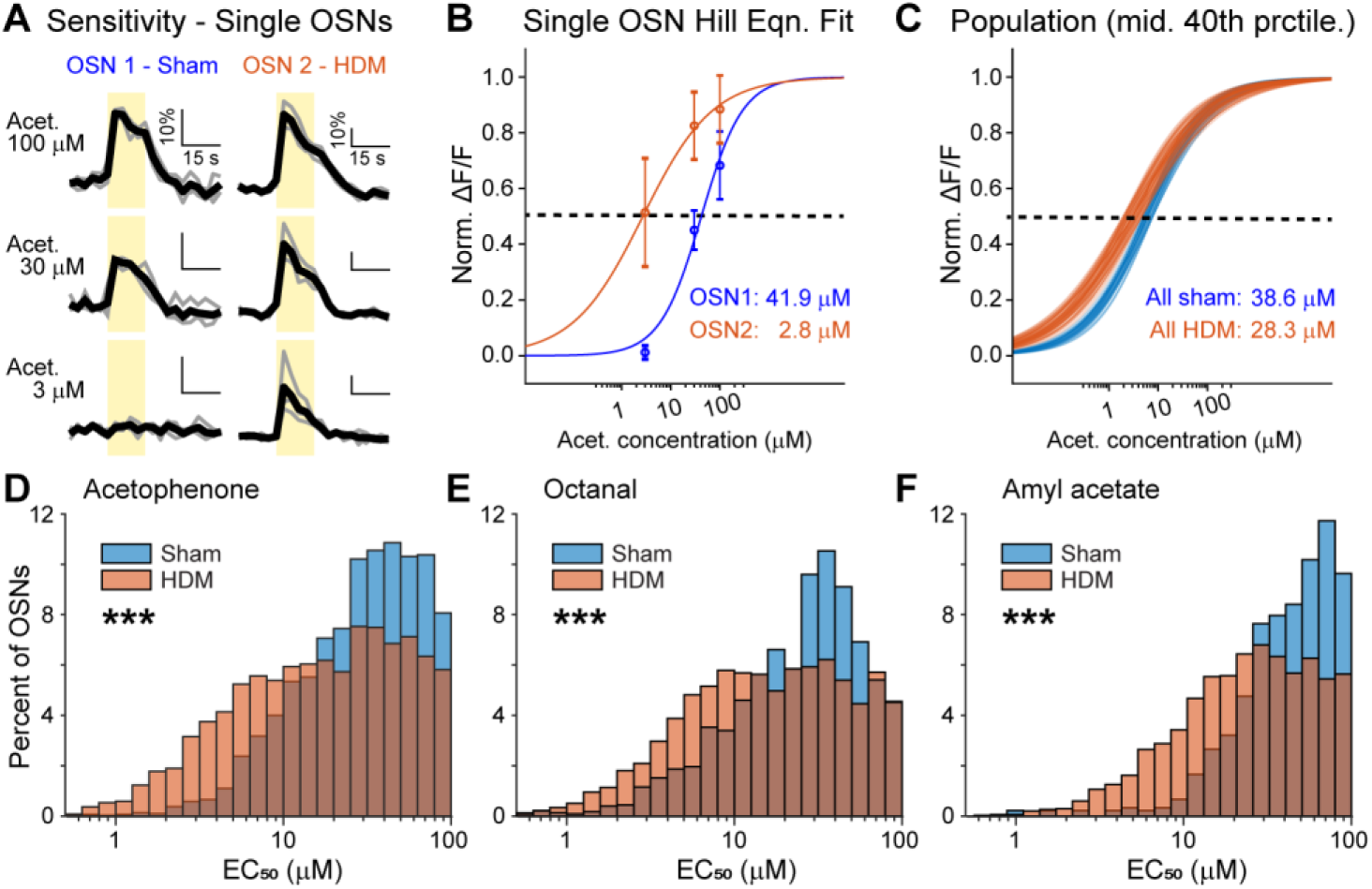
OSN odorant sensitivity increases following HDM treatment. (**A**) Example across-trial GCaMP6s Ca^2+^ responses to stimulation with acetophenone at 3, 30, and 100 μM. Gray traces show the first 2 of 4 trials, bold black traces show the mean response. (**B**) Example best-fit solutions to the Hill Equation for representative OSN responses to acetophenone. Open circles show the across-trial mean response, and the error bars reflect standard error of the mean (n = 4 trials). OSN1 and OSN2 match the neurons in Panel A. Text indicates the EC_50_ value for each OSN Hill Equation best fit solution. (**C**) Hill Equation best fit solutions for the middle 40^th^ percentile of OSNs (30^th^ – 70^th^ percentiles). Text indicates the median EC_50_ value for sham (blue) and HDM (orange) treatment groups. (**D-F**) Histograms of EC_50_ values to acetophenone (**D**), octanal (**E**), and amyl acetate (**F**). *** Denotes p < 0.001, Wilcoxon rank sum test. Effect sizes and bootstrap 95% confidence intervals (n = 1,000 bootstraps) are as follows: (**D**) Hodges-Lehmann estimator (median difference between groups): 10.23 [9.01 – 11.32], (**E**) Hodges-Lehmann estimator: 6.32 [5.16 – 7.50], (**F**) Hodges-Lehmann estimator: 18.11 [15.6899, 20.6547].

### HDM decreases OSN sensory adaptation

OSN sensitivity is related to the threshold for odorant detection, but OSNs are known to adapt their odorant responsiveness over time, most frequently a reduction in the responsiveness to repeated odorant exposure compared the initial conditions (“sensory adaptation”)^49, 50^. Given that OMP expression was upregulated in HDM-treated MOE tissues (Fig. 4), and that OMP can serve as a cAMP buffer^36^, we proceeded to quantify across-trial sensory adaptation from these same OCPI Ca^2+^ imaging data. We quantified across-trial (n = 4 trials) sensory adaptation of all strongly odorant-responsive OSNs for all concentrations, then compared the adaptation characteristics of OSNs in HDM-exposed tissues to sham controls (Fig. 7).

**Figure 7.**
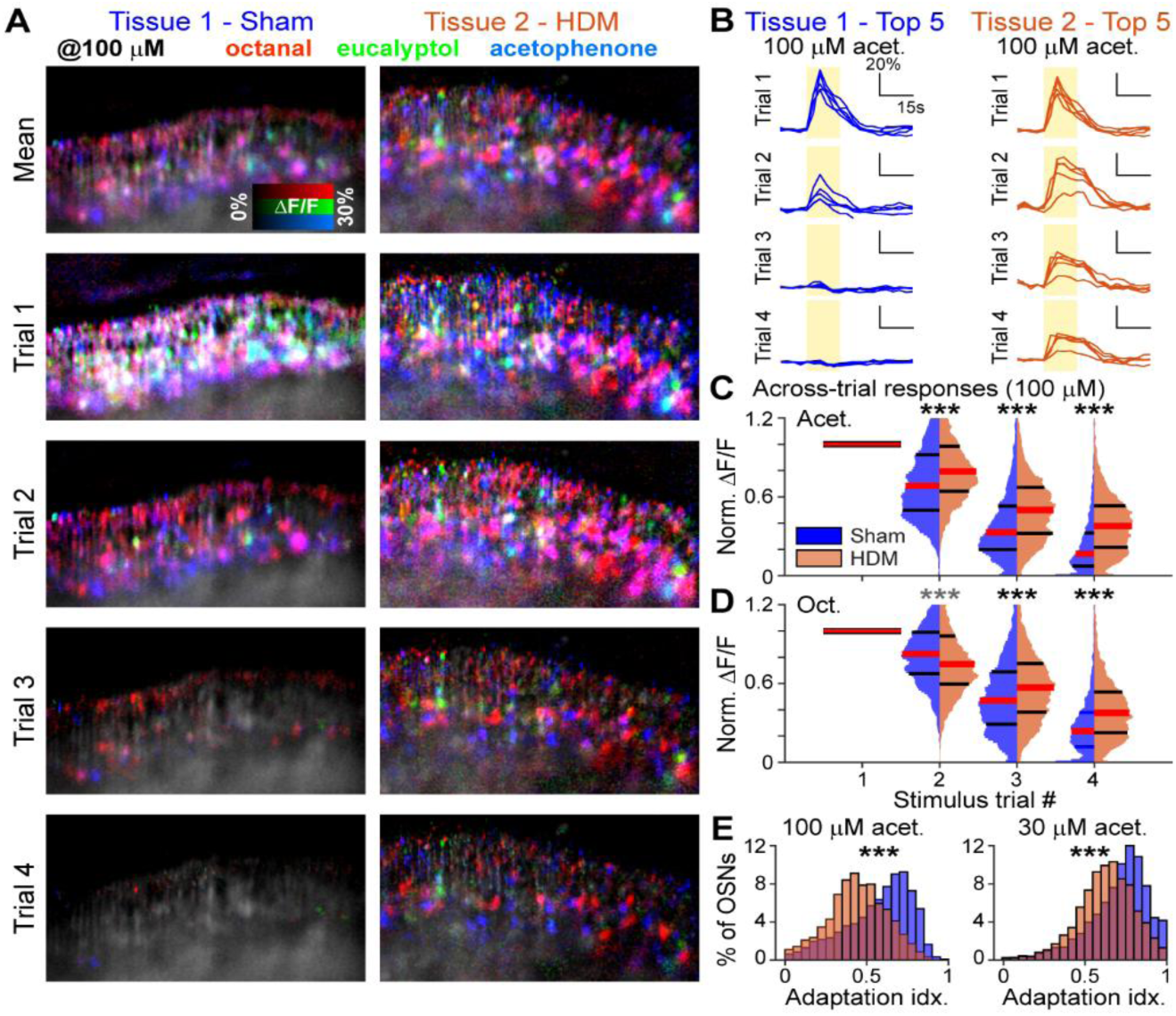
OSNs from HDM-treated MOE samples demonstrate reduced across-trial stimulus adaptation. (**A**) Pseudocolored ΔF/F image of representative MOE tissues assessed via OCPI microscopy. Mean (top row) and single-trial ΔF/F images (rows 2-5) show stimulus adaptation in Sham (left column) and HDM-treated (right column) samples to 100 μM octanal (red), eucalyptol (green) and acetophenone (blue). Pixels with other colors (yellow, magenta, white, etc.) indicate responses to multiple odorants at this concentration. (**B**) Across-trial responses of the 5 OSNs demonstrating the strongest Trial 1 responses to 100 μM acetophenone from Tissue 1 (sham, left) or Tissue 2 (HDM-treated, right) from Panel A. (**C**) Normalized across-trial ΔF/F response distributions for all 100 μM acetophenone responsive cells from Sham (blue) and HDM-treated (orange) samples. Red horizontal bars indicate group medians. Black horizontal bars indicate the 25^th^ and 75^th^ percentiles. *** indicates p < 0.001 (Wilcoxon rank sum test, Sham vs. HDM-treatment groups). Hodges-Lehmann effect size estimates and boostrap confidence intervals (n = 1000 bootstraps) were as follows: Trial 2: 11.2% [9.9 – 12.4%], Trial 3: 13.9% [12.8 – 14.9%], Trial 4: 17.2% [16.3 – 18.0%]. (**D**) Same as in Panel C, but for OSNs responsive to 100 μM octanal. Gray *** indicate the effect direction for Trial 2 responses to 100 μM octanal were slightly increased, rather than decreased, at this combination of stimulus, concentration, and trial (Hodges-Lehmann Estimator: 6.2% [5.0 – 7.3%] facilitation). Other estimates and 95% confidence intervals for adaptation were: Trial 3: 8.2% [6.9 – 9.6%], Trial 4: 12.9% [11.9 – 13.8%]. (**E**). Stimulus adaptation index (cumulative adaptation across Trials 2-4) for acetophenone at 100 μM (left) and 30 μM (right). *** indicates p < 0.001 (Wilcoxon rank sum test). Hodges-Lehmann effect size estimates and 95% confidence intervals were: 100 μM acetophenone: 14.8% [14.0 – 15.8%], 30 μM: 8.5% [7.8 – 9.2%].

OSNs in both HDM-exposed and sham tissues demonstrated across-trial stimulus adaptation (Fig. 7A-B). However, OSNs in HDM-exposed tissues showed decreased levels of stimulus adaptation compared to sham controls (Fig. 7B-C). For example, in response to the maximum concentration of acetophenone (100 μM), we observed a systematic reduction in across-trial sensory adaptation in HDM-treated tissues, resulting in a higher fraction of OSNs that remained robustly responsive at Trial 4 (Fig. 7C). The same overall result was observed, to a slightly reduced degree, in 100 µM octanal-responsive OSNs (Fig. 7D), suggesting that reduced sensory adaptation in HDM-treated tissues is not restricted to an individual odorant. Additionally, comparing the cumulative amount of adaptation (encompassing Trials 2-4) demonstrated by OSNs between HDM-exposed and sham tissues revealed shifts towards lower amounts of stimulus adaptation across odorant concentrations, suggesting that adaptation findings are not restricted to only the strongest stimulation conditions (Fig. 7E). Cumulatively, these Ca^2+^ imaging results suggest that acute environmental exposure to HDM aeroallergen sensitizes the main olfactory epithelium to odorants, and limits stimulus adaptation, each of which preserve or increase OSN stimulus-responsiveness.

## Discussion

These studies evaluated the effects of acute HDM exposure on the immune cell populations and OSN physiology in the MOE with the goal of better understanding how these populations maintain the nasal mucosa in the face of environmental challenges. Our results indicated that acute allergen exposure over the course of 7 days causes subtle changes to the distribution of subsets of immune cell populations and modulates OSNs physiology. Live *ex vivo* MOE Ca^2+^ imaging demonstrated an increase in OSN odorant sensitivity and a decrease in adaptation following HDM exposure. These results suggest that acute environmental exposure to potent aeroallergens sensitizes the MOE to odorants without inducing a parallel shift in immune cell phenotypes.

Immune cell phenotyping of the MOE indicated small changes following HDM exposure (Fig. 2). This was somewhat surprising, given the expected immunogenicity of HDM extracts, which include potent immunostimulatory molecules like lipopolysaccharide^51–53^.We did observe small increases, though, in MOE DCs and NK cells. The upregulation of these populations could be indicative of an innate immune response^37, 54^. DCs are antigen presenting cells, and their upregulation may indicate a role for MOE DCs in antigen surveillance. Macrophages and monocytes have similarities to DCs in that they express pattern-recognition receptors as a part of the mononuclear phagocyte population^55, 56^, but we did not observe changes in the abundance of macrophages or monocytes in the HDM-treated MOE. These cells are often responsive to signaling molecules like alarmins and cytokines, and this lack of change may indicate that these signaling molecules are not upregulated by acute HDM exposure. To induce more robust inflammatory response, engaging both the innate and adaptive sides of the immune response may be needed. Future studies may consider a chronic exposure model (*i.e.* weeks of exposure) in order to drive more robust inflammatory changes and alter immune cell populations in the MOE. A chronic exposure and sensitization model might induce an adaptive immune response, including HDM specific T and B cells^4, 6, 37^, and perhaps an increase in type 2 (*i.e.* allergic) cytokines such as IL-4, IL-5, and IL-13. Another limitation of this approach is that the immune cell populations in the MOE constitute only 3-4% of the total cells in this tissue^32, 34, 58^. This may contribute to between-sample variability, obscuring potential changes beneath the limit of detection. Overall, the mild changes we observed in response to HDM treatment indicates that this paradigm, despite introducing a broad spectrum of immunogenic molecules to the MOE, does not induce a widespread inflammatory response in this tissue.

An important consideration in this paradigm is the potential for regional variability in the distribution of immune cell types in the MOE. Flow cytometry indicated changes in the abundance of immune cells in the anterior/dorsal versus posterior-lateral MOE (Fig. 3). The anterior-dorsal MOE had overall higher levels of CD45+ cells, and higher levels of B cells, but a lower proportion of eosinophils compared to the posterior/lateral MOE. These differences may reflect specialization of neuro-immune functions in different MOE subregions. Multiple lines of evidence show regional specialization of function in the MOE, from receptor distributions^27–29, 59^, olfactory bulb innervation^60, 61^, and access to airborne molecules^62, 63^. Future studies will be necessary to further investigate the region-specific responses of MOE immune cell types to environmental challenges.

In contrast to the relatively minor effects of HDM treatment on immune cells, we saw a robust response of OSNs to HDM treatment (Figs. 4-7). OMP expression was upregulated in the lamina propria of the MOE (Fig. 4). This change was most observed in axon bundles in the lamina propria, which extend through the cribriform plate into the olfactory bulb, where the OSN axons terminate in MOB glomeruli^54, 64^. Upregulation of OMP in the axon bundles has been noted in other studies of olfactory manipulations, suggesting that our HDM exposure model modifies OSNs similar to naris occlusion and other challenges^40–42^. OMP has long been associated with mature OSNs but has also been associated with olfactory signal transduction and physiology^36, 65, 66^.

We directly measured OSN activity on the anterior/dorsal region of the MOE to assess potential changes to OSN physiology. We observed increases in odorant sensitivity and reduced sensory adaptation following HDM exposure compared to sham controls. Direct tests of OSN function in the native epithelium are rare^47, 67–69^, making it challenging to directly compare these findings to other studies. The olfactory stimuli used were not chosen because they were familiar to the animal, but because they were well-studied monomolecular odorants that had been presented at similar concentration ranges in other studies^70^. These specific stimuli and concentrations were highly effective at activating OSNs in the anterior/dorsal MOE region, but because of zonal differences in OR distribution, these stimuli may not represent the effects of HDM treatment on all OSNs. Changes in odor sensitivity can be affected by alterations in GPCRs or other components along the signal transduction cascade^71, 72^. Following our acute allergen model to HDM, we did not observe loss of OSNs but did observe increases in odorant sensitivity. Taken together with the increased axonal OMP, these results indicate that increased OMP happens in parallel with increased capacity to detect odorants following aeroallergen exposure. Future studies would be necessary to investigate downstream implications of these results, either at the level of the olfactory bulb glomeruli, downstream neurons, or behavior^44^.

A limitation of the OCPI imaging paradigm for MOE investigation is that it requires tissue submersion, and aqueous rather than airborne odorant stimulus delivery. Even still, our results suggest a somewhat unexpected increase in odorant sensitivity following HDM treatment, along with reduced sensory adaptation. Each of these features suggest OSNs in HDM-treated MOE tissues engage cellular processes that enhance chemosensory signal transduction. It is noteworthy that HDM treatment causes similar upregulation and sensitization of OSNs as naris occlusion^38, 39^, which restrict access of the nasal mucosa to external odorants. It may be the case that HDM treatment causes a thickening of the mucosal layer that – similar to naris occlusion – prevents odorant access to OSN dendrites. Our immunohistochemical procedures, and live imaging protocols, both wash away mucus, and thus not able to detect such a change. Importantly, these experiments cannot determine whether the changes observed in OSNs come in parallel with behavioral enhancement of odorant sensitivity and/or adaptation, which will require further investigation. Other neighboring populations of cells such as sustentacular cells, which play supportive roles in OSN physiology, may contribute to the OSN physiological changes we observed. Sustentacular cells may provide an initial response to HDM exposure, and that continued exposure might cause them to take on a more immune-like phenotype.

To our knowledge, this is the only experimental paradigm that has used an acute intranasal HDM exposure to show effects on OSN function. It is also one of the first studies to perform detailed immunophenotyping of immune cell populations in the MOE using flow cytometry. Our work alongside others, reveals that the MOE responds to environmental aeroallergen exposure by modulating aspects of OSN signal transduction, supporting an increased odorant sensitivity and reduced adaptation to odors. In addition, our data is consistent with the current evidence that changes in OMP expression parallel alterations to OSN function^36, 44^. Our work suggests that acute HDM exposure does not cause a massive inflammatory response in immune cells but does elicit an overall tissue response that preserves OSN function. These results do not reveal any major reorganization of neuro-immune communication in the acute HDM exposure paradigm, but also leave open the possibility that such crosstalk occurs, for example by changes in immune cell function that are not captured by our flow cytometry-based phenotyping. Overall, this work provides a foundation for future studies aimed at understanding how OSN-immune cell crosstalk changes across a spectrum of immune challenges and inflammatory states.

## Acknowledgements

We thank members of the Rowe and Meeks Laboratories for technical support, guidance, and feedback. This work was supported by core facility resources via the University of Rochester Center for Advanced Research Technologies, specifically the Center for Advanced Light Microscopy and Nanoscopy and (Emma Norris, Julie Zhang, and V. Kaye Thomas) and the Flow Cytometry Shared Resource (Matthew Cochran and Steven Polter).

