## Supplementary Information for "Effects of acute intranasal allergen exposure on resident immune cells and sensory neurons in the mouse olfactory epithelium"

**Supplementary Table 1. Spectral flow cytometry antibodies**

| **Spectral Flow Cytometry Panel** | | | | | |
|---|---|---|---|---|---|
| **Antigen** | **Fluor** | **Dilution** | **Clone** | **Catalog #** | **Supplier** |
| Viability Stain | Live Dead Blue | 1:1000 | n/a | L23105 | Thermo Fisher |
| CD45 | BUV395 | 1:200 | Clone: 30 F11 | 564279 | BD |
| MHC II | APC-Cy7 | 1:1000 | M5/114.15.2 | 107628 | Biolegend |
| NK1.1 | BV510 | 1:100 | PK136 | 108737 | Biolegend |
| F4/80 | AF488 | 1:100 | BM8 | 123120 | Biolegend |
| CD11b | BV711 | 1:400 | M1/70 | 101242 | Biolegend |
| CD4 | PE Cy7 | 1:400 | GK1.5 | 100422 | Biolegend |
| CD8a | BV650 | 1:100 | 53-6.7 | 100742 | Biolegend |
| Ly6G | AF647 | 1:200 | 1A8 | 120610 | Biolegend |
| Ly6C | BV421 | 1:200 | HK1.4 | 128032 | Biolegend |
| CD3 | BUV805 | 1:50 | 17A2 | 741982 | Biolegend |
| CD11c | BV786 | 1:100 | ZM3.8 | 563735 | BD |
| Siglec-F | PE-CF594 | 1:100 | Clone E50-2440 | 562757 | BD |
| CD19 | BV605 | 1:100 | 6D5 | 115539 | Biolegend |
| Cx3cr1 | BB700 | 1:50 | Z8-50 | 567812 | BD |

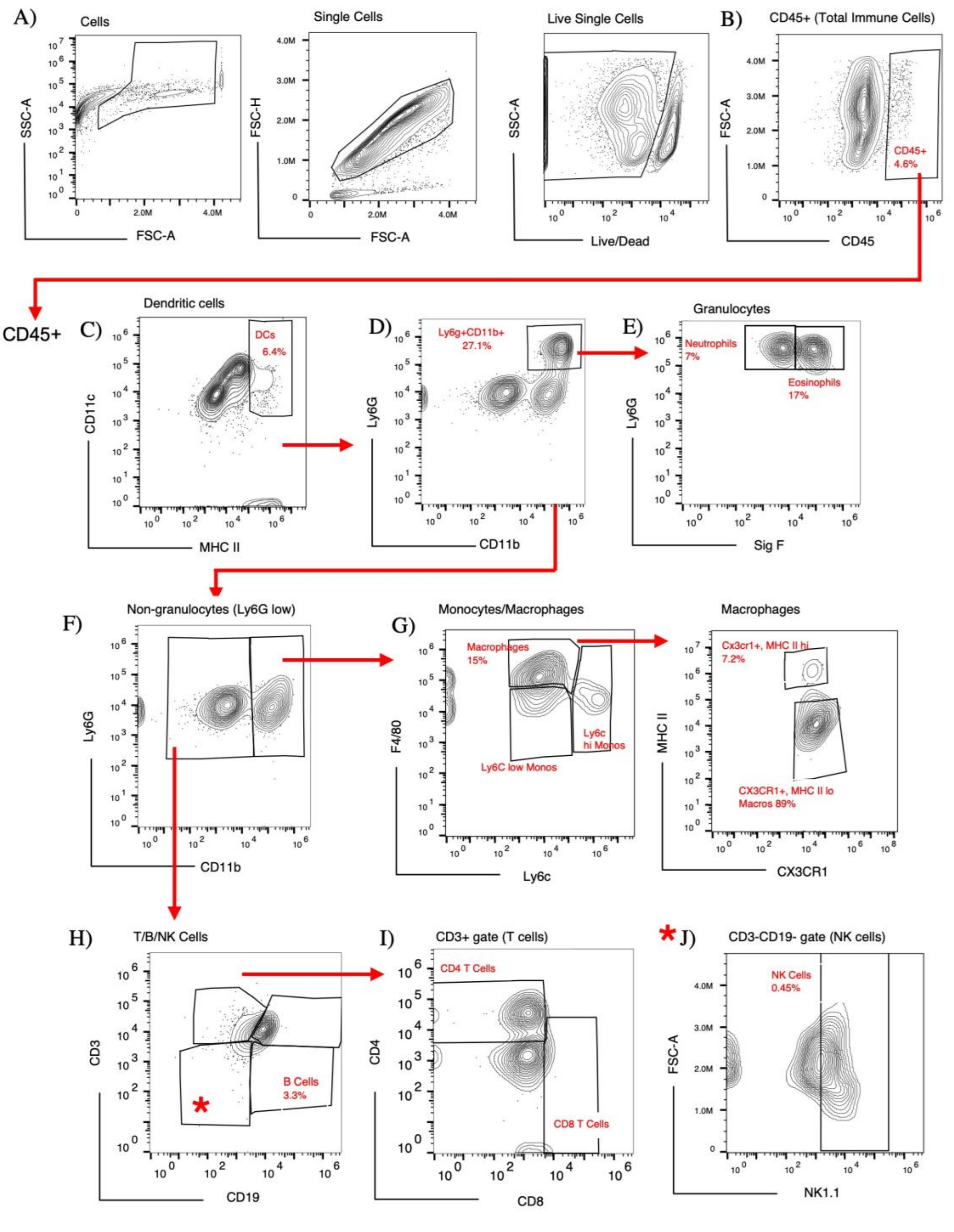


**Supplementary Figure 1. A schematic overview of the multi-step gating strategy to identify immune cell populations in the MOE.** Example is from a female (age 6-8 weeks) C57BL/6J mouse MOE tissue. **(A)** Cells were determined based on events in SSC-A vs FSC-A to discriminate from debris. To discriminate doublets, events in FSC-A vs. FSC-Height (FSC-H) where considered as singlets. Live single events were further identified as live-dead blue low/negative. **(B)** $CD45^{+}$ hematopoietic cells were determined from the live-single events. The CD45+ gate was then used for downstream gating of the following immune cell types: **(C)** dendritic cells (DCs), **(D)** Non-DC events were then used to identify granulocytes that were Ly6G+, CD11b+ and subsequently gated into **(E)** neutrophils and eosinophils. **(F)** Non-granulocytes. CD11b positive and negative populations were used to identify **(G)** Monocytes and macrophages, **(H)** B cells were identified as CD3-, CD19+ cells, and further gating for **(I)** T cell and **(J)** NK cell populations.

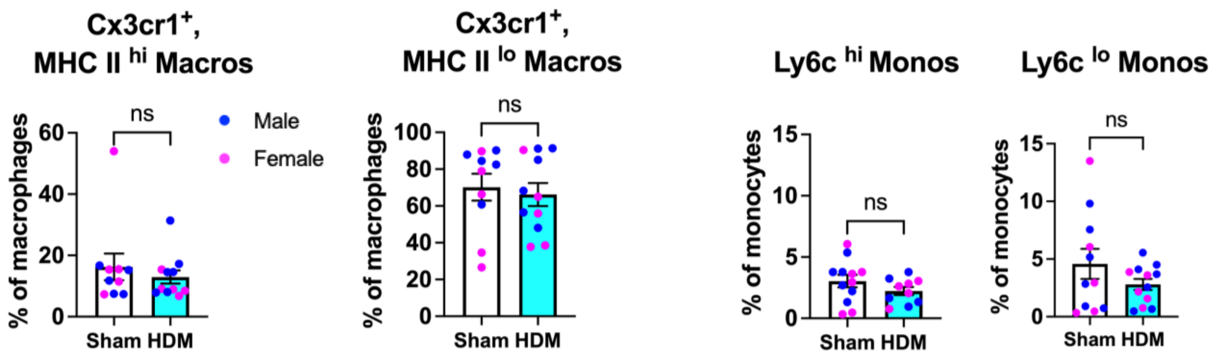


**Supplementary Figure 2. Investigation of MOE macrophages and monocytes.** Monocyte and macrophages were gated into subpopulations based on expression levels of MHC II, Cx3cr1, and Ly6c to denote inflammatory immune populations and surveillance cells. In each graph, magenta symbols indicate results for females, and dark blue symbols indicate results for males. ns: not significant.

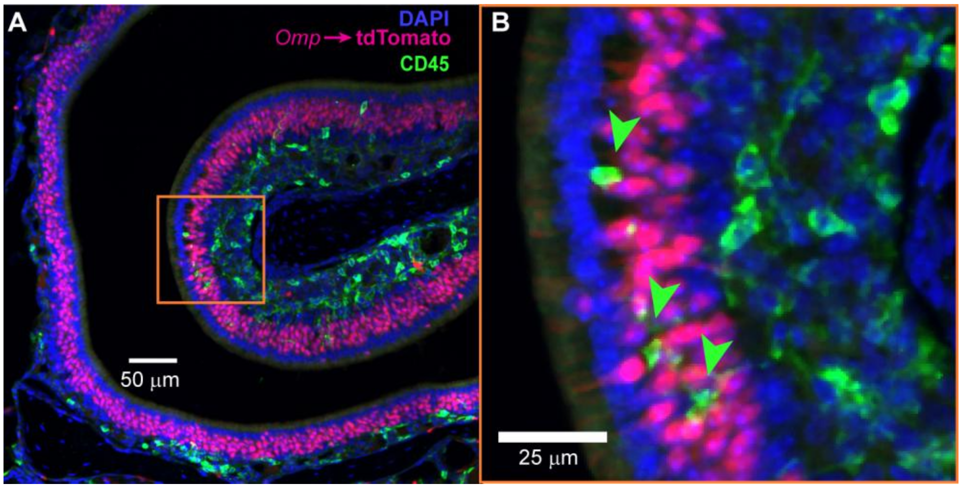


**Supplementary Figure 3**: **Representative confocal image of CD45$^{+}$ immune cells spatially localized to OSNs in the MOE.** Omp-cre x Ai9 (Cre-dependent cytoplasmic tdTomato) transgenic mice were used to identify mature OSNs (via tdTomato expression, magenta). Immunostaining against CD45 identified immune cells, and DAPI staining identified nuclei. Image captured by confocal microscopy at (A) 10X and (B) 20X magnification.

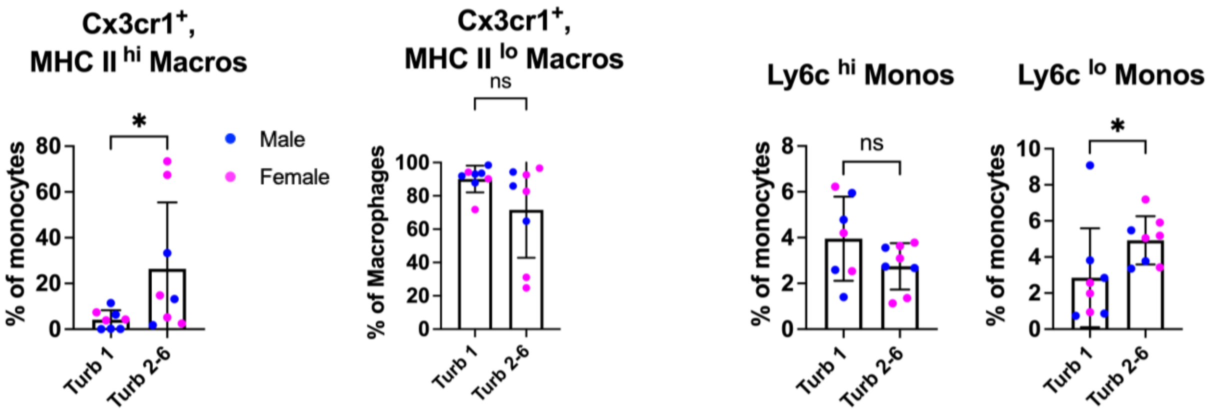


**Supplementary Figure 4. Investigation of macrophages and monocyte distributions across different MOE regions.** Identification of each subset was performed as in Supplementary Figure 2. In each graph, magenta symbols indicate results for females, and dark blue symbols indicate results for males. * $p < 0.05$ (Mann-Whitney test); ns: not significant.
